# Electrophysiological correlates of the flexible allocation of visual working memory resources

**DOI:** 10.1101/746164

**Authors:** Christine Salahub, Holly A. Lockhart, Blaire Dube, Naseem Al-Aidroos, Stephen M. Emrich

## Abstract

Visual working memory is a brief, capacity-limited store of visual information that is involved in a large number of cognitive functions. To guide one’s behavior effectively, one must efficiently allocate these limited memory resources across memory items. Previous research has suggested that items are either stored in memory or completely blocked from memory access. However, recent behavioral work proposes that memory resources can be flexibly split across items based on their level of task importance. Here, we investigated the electrophysiological correlates of flexible resource allocation by manipulating the distribution of resources amongst systematically lateralized memory items. We examined the contralateral delay activity (CDA), a waveform typically associated with the number of items held in memory. Across three experiments, we found that, in addition to memory load, the CDA flexibly tracks memory resource allocation. This allocation occurred as early as attentional selection, as indicated by the N2pc. Additionally, CDA amplitude was better-described when fit with a continuous model predicted by load and resources together than when fit with either alone. Our findings show that electrophysiological markers of attentional selection and memory maintenance not only track memory load, but also the proportion of memory resources those items receive.

---

More likely than not, it is much easier for you to recall the names of the characters from the last television show that you watched than what you were wearing while you watched it. This bias in memory is in part due to the fact that what we allocate more attention to is remembered with greater detail^1^. Indeed, numerous studies of long-term memory have established that attention prioritizes relevant information to be encoded into memory^2–4^. Attention also affects the maintenance and quality of information stored over shorter periods of time, such as in visual working memory^5–7^ (VWM). In fact, given that VWM is limited in capacity, several models of VWM have suggested that attention may play a critical role in determining what information gains access to these finite storage resources^8–11^.

One potential mechanism through which attention may drive working memory performance is by filtering out irrelevant distractors^12^. Filtering efficiency has been quantified using measurements of electrophysiological brain activity related to working memory storage – specifically an event-related potential (ERP) called the contralateral delay activity^13–16^ (CDA). CDA amplitude increases with the number of items stored in VWM, saturating as memory load increases beyond a few items^17^. Interestingly, when distractors are presented alongside targets in a memory display, lower-capacity individuals exhibit larger CDA amplitudes than those with higher capacities, reflecting the fact that they have encoded and stored more distractors in memory^16^. This finding has been taken as evidence that poor filtering efficiency, resulting in unnecessary storage, is a critical determinant of VWM capacity.

Several recent studies have demonstrated that it is also possible to bias attentional resources toward and away from certain items in a flexible manner, independent of the need to filter out irrelevant distractor items. This bias can be induced by associating certain stimuli with monetary incentives, or by simply varying instructions indicating the probability that an item will be probed on any given trial^7^. In this way, the proportion of attentional resources allocated to any given memory item can continuously vary anywhere between 0 and 100%. In these past studies^18,19^, it was found that working memory performance (i.e., raw error = 1/precision) was best predicted by the likelihood that an item would be probed on a given trial, independent of the overall memory load. Importantly, this relationship between probe likelihood and memory precision, which followed a power-law, could also account for changes in performance across loads; for example, the change in precision from one to two items was consistent with each item only receiving half as many attentional resources. This framework suggests that attention does more than just restrict or grant access to VWM; rather, it also flexibly distributes resources amongst objects based on their respective priorities. In other words, how attention is allocated between targets is as important to memory performance as whether or not it is allocated to distractors^20,21^.

If attention can be flexibly allocated across items in VWM, how might this be reflected by neural measures of attention and VWM maintenance? There is evidence that the CDA is well-described by a saturation model, which predicts a continuous increase in CDA amplitude that saturates as set size becomes larger, instead of increasing discretely and plateauing at memory capacity^17^. This finding suggests that the CDA, much like VWM performance, may be more flexibly affected by memory load than previously thought. Yet, it is currently unknown whether the CDA is also flexibly affected by the prioritization of memory items instead of, or in addition to, changes in memory load.

Prioritization could also be tracked by ERP components that precede memory maintenance, such as attentional selection and suppression. That is, one way that flexible prioritization could be accomplished is through the specific up-weighting of goal-relevant over irrelevant information (as opposed to down-weighting of goal-irrelevant information). Attentional selection can be tracked by the N2pc, a lateralized component which specifically reflects the enhancement of an item^22–24^. Alternatively, it could be that prioritization is accomplished through the active suppression or down-weighting of goal irrelevant information. This can be measured by the distractor positivity (P_D_): a lateralized component that is observed when distractors are presented laterally in the stimulus display^25–28^. These two components can thus be used to disentangle the underlying mechanisms of prioritization: whether through selective enhancement of relevant information (N2pc) or suppression of irrelevant information (P_D_).

Consequently, to determine the effect of resource allocation on the CDA, as well as whether prioritization is driven by selective enhancement of high-priority items or inhibition of low-priority items, we conducted three experiments in which the allocation of memory resources across items was manipulated in a continuous-report delayed-recall task. For these experiments, we use the term *memory load* to refer to the number of items with greater than zero percent likelihood of being probed. We use the terms *resource allocation* or *probability* to refer to the likelihood that one item, or a set of items, will be probed. In Experiment 1, we examined how changes in resource allocation amongst memory items influenced the CDA in comparison to the typical effect of memory load. To do this, participants were asked to remember the colors of four laterally presented items that were either equally likely to be probed, or where a spatial cue indicated that one item was more likely to be probed than the others (i.e. 50% or 100% likely to be probed). Thus, while participants should always be allocating 100% of memory resources to all four items, how the resources were distributed across items varied. If the CDA is unaffected by resource allocation, and is only affected by memory load, then CDA amplitude should only reflect the total number of items to be stored, regardless of attentional priority.

In Experiments 2 and 3, we took advantage of an attribute of the CDA, N2pc, and P_D_ – that these components are only sensitive to laterally presented stimuli and not stimuli presented on the vertical midline – to separately manipulate the effects of memory load and resource allocation. In Experiment 2, two items were presented laterally and two vertically, and a featural cue indicated whether the lateral or vertical items were more likely to be probed. Thus, this design allowed us to manipulate the proportion of memory resources specifically allocated to lateral items. If these ERP components reflect strategic resource allocation, then they should increase in amplitude with increasing probe likelihood (i.e., greater resource allocation). However, if these components are only affected by overall load, then item prioritization should have no effect.

In Experiment 3, we tested whether the CDA reflects the allocation of memory resources even in the absence of prioritization cues. To do so, we manipulated the total number of to-be-remembered items (four or six), while systematically changing the number of items presented laterally. In this way, we could simultaneously manipulate lateral memory load and proportion of memory resources allocated to the lateral items. We predicted that if the CDA tracks resource distribution across the lateral and vertical items, then CDA amplitude should reflect the proportion of total memory resources allocated to lateral items in addition to lateral memory load.

## Results

### Behavioral: Experiments 1 - 3

Because we manipulated proportion of memory resources per item across all three experiments, and were interested in how behavior changed as a function of resource allocation, behavioural results were collapsed across experiments.

To compare how performance changed as a function of resource allocation, all data points were fit to a power-law function. Consistent with past findings^18,19^, this provided a good fit (Fig. 1), with the model accounting for around 79% of the variance in the data, adjusted-R^2^ = .79, *RMSE* = 6.26. These results demonstrate that the proportion of memory resources allocated to an individual item was highly predictive of behavioral precision for that item. Moreover, percent of memory resources allocated to an item better predicted behavioral precision than memory load alone: adjusted-R^2^ = .35, *RMSE* = 10.97, Δ*BIC* = 9.71. Thus, regardless of the behavioural manipulation (i.e., spatial cues; feature-based cues; lateralized memory load), precision is strongly predicted by the percentage of resources allocated to the probed item.

**Figure 1.**
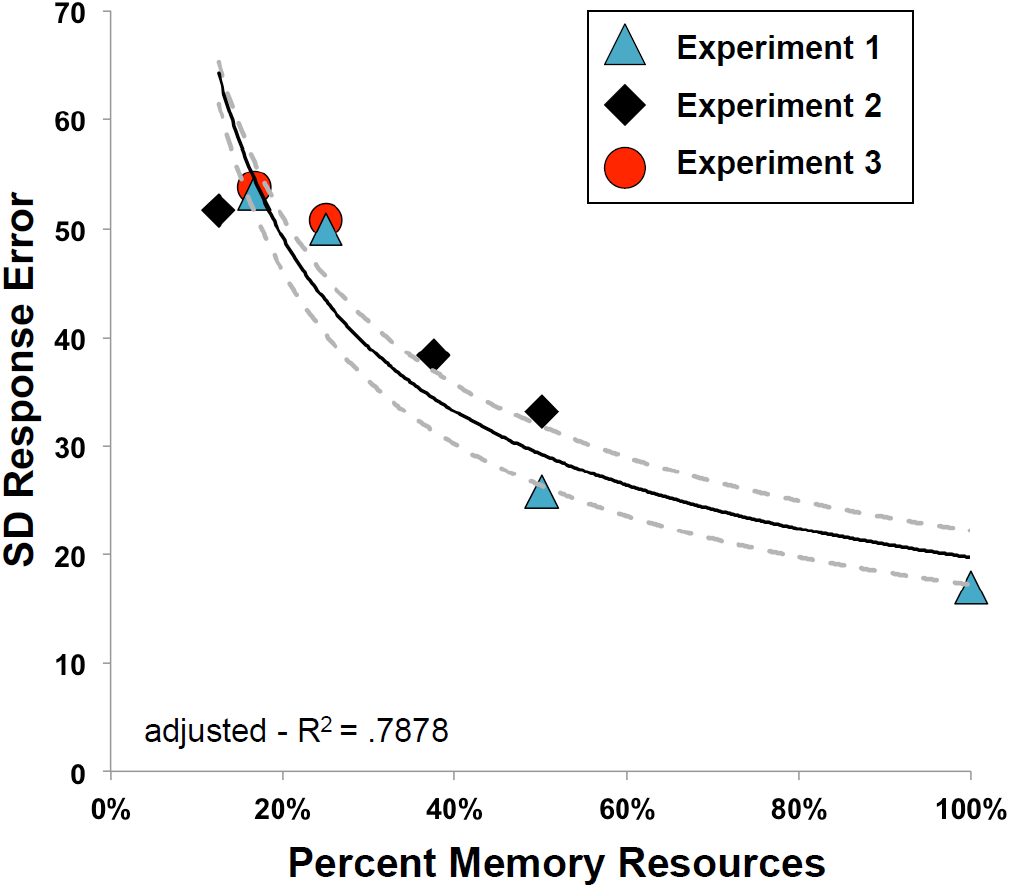
Standard deviation of raw response error by percent memory resources in each experiment, fit with a power-law. Dashed grey lines represent fits performed on the 95% confidence interval of the condition means.

### ERPs

#### Experiment 1

In this experiment, we sought to examine how the prioritization of one item over the others affected the CDA. Four lateral memory items were always presented, and spatial cues indicated the number of items to be remembered, as well as the likelihood of a given item to be probed (Fig. 2A). Based on past demonstrations that the CDA primarily reflects VWM load, we expected there to be a larger CDA amplitude in the 4-Cues/100%-Valid condition than the 1-Cue/100%-Valid condition; remembering four items results in a larger CDA than remembering a single item. What remains an open question, however, is how the CDA changes when resources are distributed unequally across the four memory items. Thus, comparing the 4-Cues/100%-Valid and the 1-Cue/100%-Valid conditions to the 1-Cue/50%-Valid condition provides an initial test of how resource allocation affects CDA amplitude.

**Figure 2.**
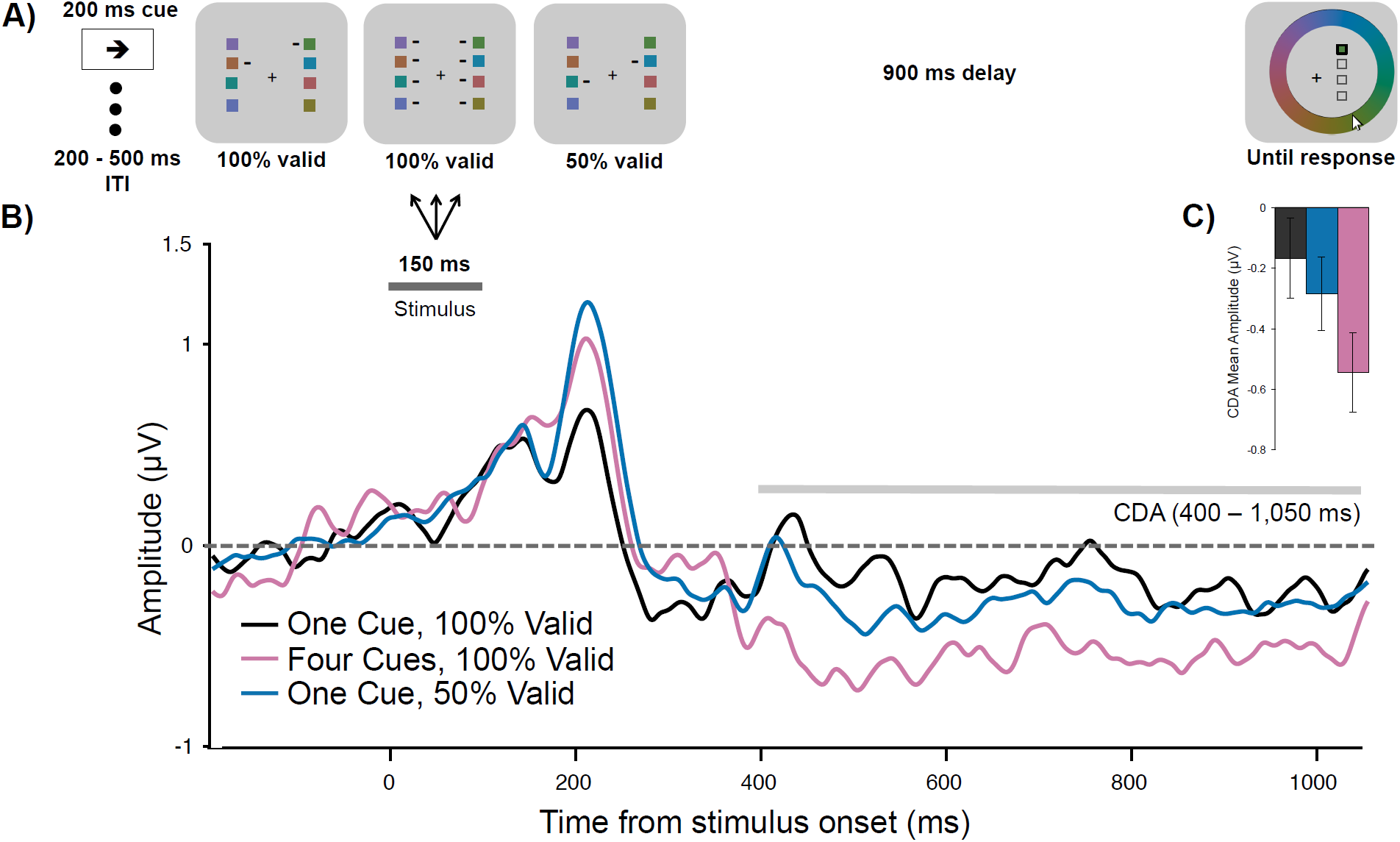
A) Task schematic of Experiment 1.B) Grand average difference waveform (N = 20) at the average of 5 posterior channel pairs, time-locked to stimulus onset. Positive is plotted up. Filtered at 30 Hz for visualization purposes only. C) Bar chart of mean CDA amplitudes in each condition. Error bars reflect within-subject 95% confidence intervals.

#### CDA

The spatial prioritization cues influenced CDA amplitude (Fig. 2B; main effect of Condition, *F*(2,38) = 5.83, *p* = .006, η^2^_p_ = .24, BF_10_ = 6.94). Consistent with a memory load effect, there was a more negative CDA amplitude when all four items were cued (*M* = −0.55 μV, *SD* = 0.66 μV), than when one item was cued at 100% validity (*M* = −0.17, *SD* = 0.70 μV), *t*(19) = 3.43, *p* = .003, d = 0.77, BF_10_ = 14.94. CDA amplitude in the 1-Cue/50%-Valid condition (*M* = −0.29 μV, *SD* = 0.52 μV) was not significantly different from the 1-Cue/100%-Valid condition, *t*(19) = 0.98, *p*_*bonf*_ = 1, d = 0.22, BF_10_ = 0.36, or from the 4-Cues/100%-Valid condition, *t*(19) = 2.38, *p*_*bonf*_ = .084, d = 0.53, BF_10_ = 2.20. Thus, though the CDA amplitude in the 1-Cue/50%-Valid condition was numerically smaller than in the 4-Cues/100%-Valid condition, this difference was not born out in the inferential statistics. Instead, the CDA amplitude in the 1-Cue/50%-Valid condition appeared to be in between the amplitudes of the other conditions (see Fig. 2C).

#### Experiment 2

In Experiment 1, we replicated the typical effect of memory load on CDA amplitude and the behavioral effect of resource allocation on memory precision. However, the effects of resource allocation on CDA amplitude were less clear, as both high and low probability items were presented together laterally, resulting in a mixed electrophysiological signal. To better isolate the effects of prioritization on CDA amplitude, in Experiment 2 we separated the items in the memory array along the horizontal and vertical midlines and used feature-based priority cues (Fig. 3A). Specifically, all memory arrays comprised two items presented laterally, and two presented on the vertical midline, with the lateral items either 100%, 75%, 25%, or 0% likely to be probed, depending on the shape of those items. Because lateralized ERP components are only sensitive to laterally presented stimuli, we could systematically manipulate the proportion of lateral memory resources and thus its effect on the N2pc, P_D_, and CDA. If these components reflect strategic resource allocation, then their amplitudes should increase continuously with item priority. However, if these components reflect memory load alone, then their amplitudes should remain stable regardless of the priority manipulation.

**Figure 3.**
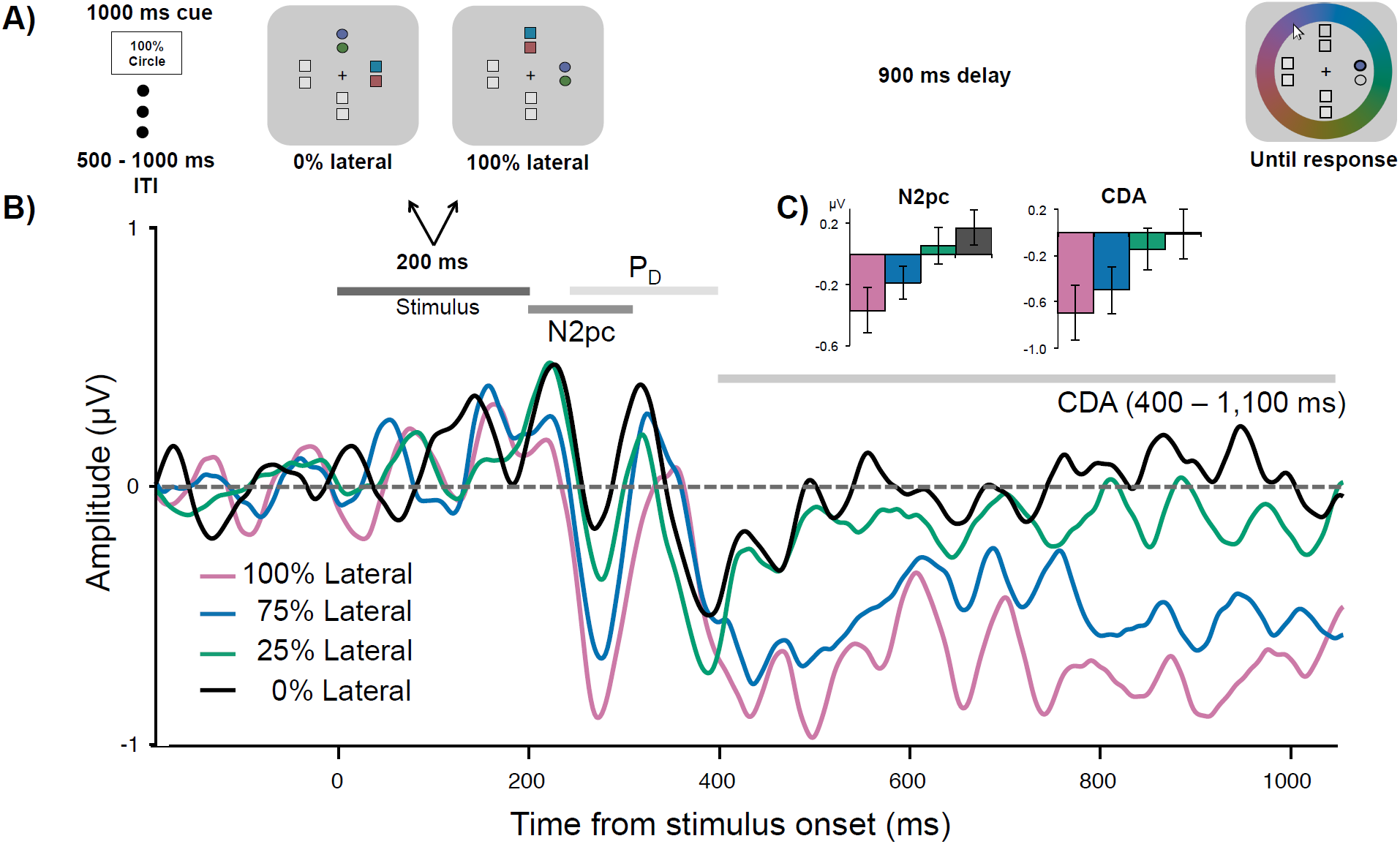
A) Task schematic of Experiment 2. In this example, it was 100% likely that the color of a circle would be probed. B) Grand average difference waveform (N = 20) at the average of 5 posterior channel pairs, time-locked to stimuli onset. Filtered at 30 Hz for visualization purposes only. C) Bar chart of mean N2pc and CDA amplitudes in each condition. Error bars reflect within-subject 95% confidence intervals.

#### N2pc

Feature-based priority influenced the amplitude of the N2pc (Fig. 3B; main effect of Condition, *F*(2.17, 41.24) = 10.85, *p* < .001, η^2^_p_ = .36, BF_10_ = 1871.28). Overall N2pc amplitude was more negative when the items were 100% (*M* = −0.37 μV, *SD* = 0.74 μV) likely to be probed than when they were 0% (*M* = 0.17 μV, *SD* = 0.58 μV), *t*(19) = 4.82, *p* < .001, d = 1.08, BF_10_ = 238.64, or 25% likely to be probed (*M* = 0.06 μV, *SD* = 0.52 μV), *t*(19) = 3.22, *p* = .005, d = 0.72, BF_10_ = 10.02. The N2pc was also larger when the lateral items were 75% likely (*M* = −0.19 μV, *SD* = 0.72 μV) compared to 0%, *t*(19) = 3.32, *p* = .004, d = 0.74, *BF*_10_ = 12.21, or 25% likely to be probed, *t*(19) = 2.57, *p* = .019, d = 0.57, BF_10_ = 3.05. There were no significant differences between N2pc amplitudes in any other condition comparison, *p*s > .069, ds < 0.44, BFs_10_ < 1.08. However, the overall N2pc amplitude did become linearly more negative as item priority increased, adjusted-R^2^ = .99, *RMSE* = 5.54, linear contrast: *t*(19) = 5.66, *p* < .001, suggesting that individuals could flexibly allocate their attention toward an item depending on how important it was to the trial (see Fig. 3C). Interestingly, fractional peak latency did not differ between conditions, *F*(1.81, 29.03) = 1.70, *p* = .202, η^2^_p_ = .10, BF_01_ = 2.29. Therefore, participants were not selecting high priority items any faster than low priority items.

#### P_D_

Permutation tests indicated that the positive area of the grand average waveform between 250 – 400 ms was not significantly different from noise in any of the conditions, 100%: *p* = .529, 75%: *p* = .30, 25%: *p* = .306, 0%: *p* = .087. Although the P_D_ was not significant, it could be that priority still had an influence on its amplitude. There was a small but non-significant effect of priority on the positive area of the P_D_, *F*(1.84, 34.96) = 2.95, *p* = .070, η^2^_p_ = .13, BF_10_ = 1.36. Therefore, there was little evidence of active attentional suppression in this task.

#### CDA

Similar to the N2pc, priority affected the amplitude of the CDA (main effect of Condition, *F*(2.02, 38.42) = 6.43, *p* = .004, η^2^_p_ = .253, BF_10_ = 85.42). More information was stored in VWM when the items were 100% likely to be probed (*M* = −0.69 μV, *SD* = 0.78 μV) than 0% (*M* = −0.02 μV, *SD* = 0.41 μV), *t*(19) = 3.34, *p* = .003, d = 0.75, BF_10_ = 12.53. Similarly, the CDA was more negative in the 100% condition than the 25% condition (*M* = −0.14 μV, *SD* = 0.36 μV), *t*(19) = 2.75, *p* = .013, d = 0.61, BF_10_ = 4.18. When the items were 75% likely to be probed, the CDA amplitude (*M* = −0.49 μV, *SD* = 0.68 μV) was more negative than in the 25%, *t*(19) = 2.15, *p* = .045, d = 0.48, BF_10_ = 1.52, and 0% conditions, *t*(19) = 2.38, *p* = .028, d = 0.53, BF_10_ = 2.20. No other comparisons were significant, *t*s < 0.18, *p*s > .18, ds < 0.31, BFs_10_ < 0.52. However, similar to the N2pc, CDA amplitude was linearly related to priority, adjusted-R^2^ = .99, *RMSE* = .03, linear contrast: *t*(19) = 4.33, *p* < .001 (see Fig. 3C). Therefore, the more likely the items were to be probed, the greater the amplitude of the CDA.

#### N2pc and CDA amplitudes predict behavioral precision

To examine whether memory resource-related changes in N2pc and CDA amplitudes predicted changes to VWM response error, a repeated-measures correlation was performed^29,30^ between mean amplitude and response error across three lateral resource conditions (25%, 75%, and 100%). It was found that attention toward the lateral shapes, as measured by the N2pc, predicted how precisely the color of the probed shape was reported, *r*_*rm*_(39) = 0.55, 95% CI = [.28, .74], *p* = < .001 (Fig. 4A). There was also a correlation between raw error and mean amplitude of the CDA, *r*_*rm*_(39) = 0.44, 95% CI = [0.15, 0.67], *p* = .004 (Fig. 4B). These findings indicate that more precise reports of the probed color were associated with larger neural responses related to attentional enhancement and memory maintenance.

**Figure 4.**
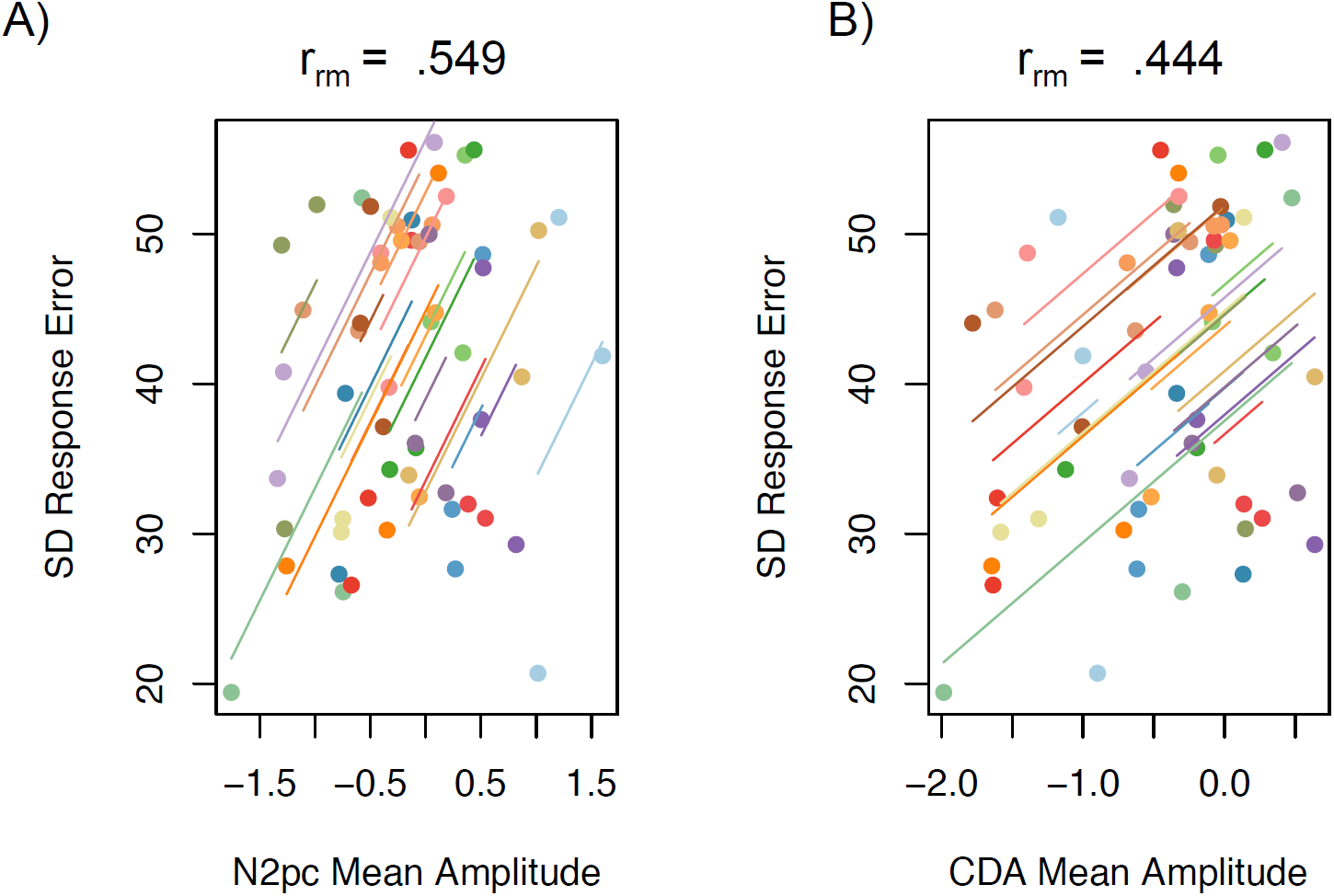
Repeated-measures correlations plots. Each colored line is the fit for three data points from each individual participant from the 100%, 75%, and 25% lateral likelihood conditions. A) Correlation between N2pc mean amplitude and standard deviation (SD) of raw response error. Lower SD indicates more precise responding. B) Correlation between CDA mean amplitude and SD of response error.

#### Experiment 3

Experiment 2 provided evidence that attentional prioritization not only affects behavioral precision in a delayed-recall task, it is also associated with a proportional increase in the amplitude of ERP components associated with attentional enhancement (N2pc) and memory maintenance (CDA). Interestingly, previous studies have found that the effect of load on behavioral precision is identical to those of prioritization; thus, splitting resources across two items results in similar memory precision as an item with 50% cue validity^18^. Consequently, to test whether the CDA similarly reflects resource allocation in the absence of prioritization cues we manipulated how many items were presented laterally, and how many vertically (Fig. 5A). There were three conditions: one item lateral and three vertical (Load 4, 25% lateral), three items lateral and one vertical (Load 4, 75% lateral), and three items lateral and three items vertical (Load 6, 50% lateral). Thus, these last two conditions had the same lateral memory load but a change in the proportion of memory resources allocated to those items. We predicted that CDA amplitude would become more negative as the proportion of lateral memory resources increased.

**Figure 5.**
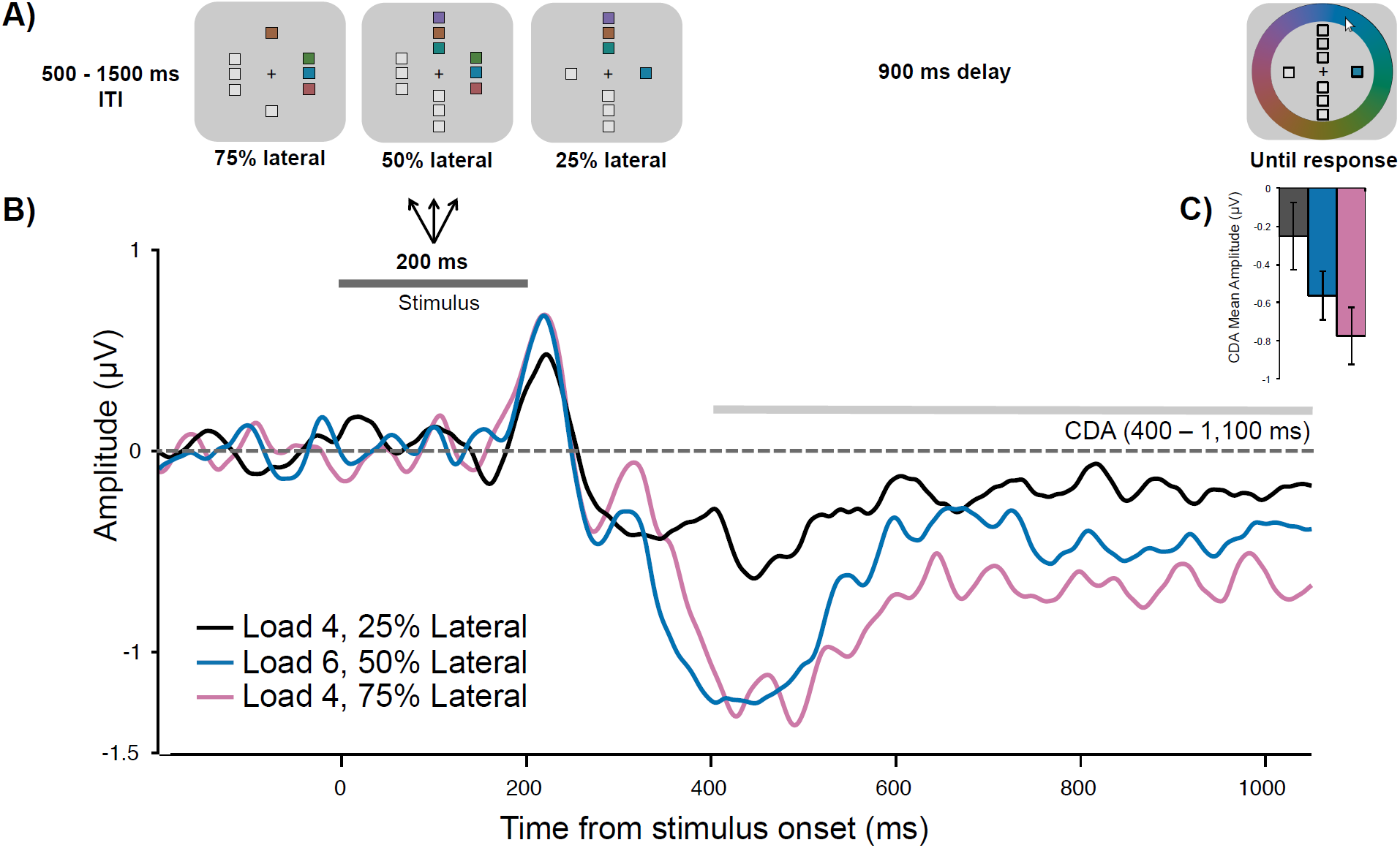
A) Task schematic of Experiment 3. B) Grand average difference waveform (N = 20) at the average of 5 posterior channel pairs, time-locked to stimuli onset. Positive is plotted up. Filtered at 30 Hz for visualization purposes only. C) Bar chart of mean CDA amplitudes in each condition. Error bars reflect within-subject 95% confidence intervals.

#### CDA

CDA amplitude was affected by Condition (Fig. 5B), *F*(2,38) = 7.60, *p* = .002, η^2^_p_ = .29, BF_10_ = 24.31, such that the amplitude was more negative when 75% of memory resources were allocated to three lateral items (*M* = −0.78 μV, *SD* = 0.64 μV) than when 25% were allocated to one lateral item (*M* = −0.25 μV, *SD* = 0.54 μV), *t*(19) = 3.4, *p* = .003, d = 0.76, BF_10_ = 14.17. When 50% of memory resources were allocated to three lateral items (*M* = −0.57 μV, *SD* = .60 μV), the CDA amplitude was more negative than when 25% of resources were allocated to one item, *t*(19) = 2.27, *p* =.035, d = 0.51, BF_10_ = 1.83. CDA amplitude in the 50% lateral condition was numerically, but not significantly, smaller than in the 75% lateral condition, *t*(19) = −1.92, *p* = .069, d = 0.43, BF_10_ = 1.08. Importantly, CDA amplitude was linearly related to the proportion of lateral resources, adjusted-R^2^ = .91, *RMSE* = .10, *t*(19) = 3.4, *p* = .009, such that amplitude became more negative as the amount of resources increased (Fig. 5C). Together, these findings suggest that even in the absence of prioritization cues, the CDA may reflect a combination of memory load *and* the amount of attention/memory resources allocated toward these items.

### CDA amplitude continuously reflects both VWM load and resource allocation

Across three experiments, the manipulation of resource allocation – whether by spatial cues, feature-based cues, or memory load – affected the amplitude of the CDA. Although these effects were sometimes small, they are consistent with previous behavioral findings (also observed here) that the magnitude of the effect on memory performance depends on the magnitude of the change in resource allocation. However, although small changes in resource allocation may only produce small effects, these effects tend to follow a predictable pattern along a continuous power-law in behavioral studies^18^. Thus, it is possible that the effect of resource allocation on ERP measures of memory maintenance should similarly follow a continuous pattern, wherein the amplitude of the CDA changes with the proportion of resources allocated to laterally presented items. It is also possible that, although resource allocation is a better predictor of memory performance than load alone, CDA amplitude may reflect a mixture of signals that combine effects of load *and* resource allocation. To examine this prediction, we tested whether the CDA amplitudes observed in Experiments 2 and 3 (which involved similar stimulus displays) were best described by one of three models: one in which CDA amplitude was predicted by load alone, another with resource allocation alone, and a model using a scaled combination of memory load and resource allocation (see Methods).

When CDA amplitudes were compared to memory load alone (Fig. 6A), the model accounted for 50% of the variance in the data, adjusted-R^*2*^ = 0.50, *RMSE* = 0.28, BIC = −17.52. However, when examining the best-fit line, the direction of this function was in the opposite direction to what was predicted, such that CDA amplitude appears not to saturate, but continues to increase with increasing memory load. This suggests that a power function does not well describe the data when fitting CDA amplitude with load, at least within the range of set sizes tested here. CDA amplitudes were even less well fit with proportion of memory resources alone (Fig. 6B), adjusted-R^*2*^ = 0.33, *RMSE* = 0.32, BIC = −15.62. However, when fitting CDA amplitude to the weighted sum of both memory load and proportion memory resources, we observed the best fit (Fig. 6C), adjusted-R^*2*^ = 0.52, *RMSE* = 0.27, BIC = −18.35. This demonstrates that the amplitude of the CDA follows a predictable continuous function that is affected both by the number of lateral items to be remembered, and by the proportion of total resources allocated to those items.

**Figure 6.**
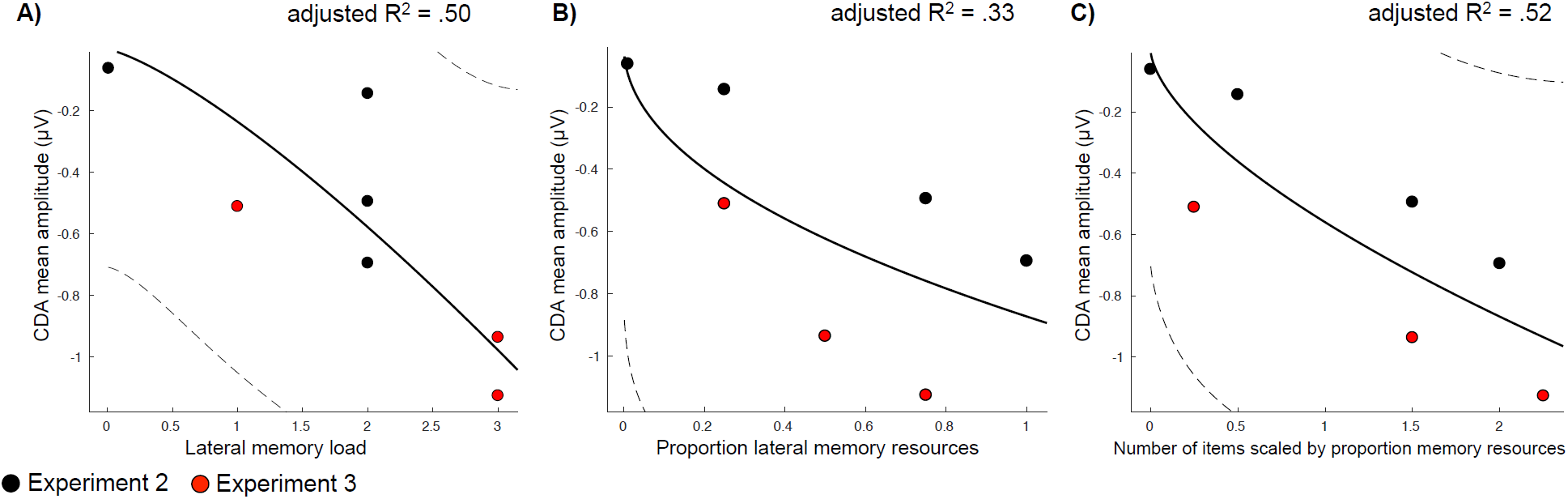
Power-law models and fits. Dotted lines represent 95% CIs of the model fit. Black dots represent condition means from Experiment 2 and red dots from Experiment 3. A) Fit between CDA mean amplitude and lateral memory load. B) Fit between CDA amplitude and proportion lateral memory resources. C) Fit between CDA amplitude and number of items scaled for both memory load and memory resources.

## General Discussion

In this study, we sought to examine the effect of prioritization on ERP markers of attentional enhancement, distractor suppression, and working memory maintenance, to better understand flexible allocation of memory resources. In Experiment 1, we found that the CDA was somewhat smaller when prioritizing one item over others than when all items were prioritized equally. In Experiment 2, we implemented a stronger manipulation of resource allocation using a systematic lateralization procedure^27,31^, demonstrating that the CDA tracked overall proportion of memory resources allotted laterally. We also found that the N2pc was reflective of priority, providing evidence that the allocation of neural resources toward to-be remembered items occurs via attentional enhancement. Moreover, both N2pc and CDA amplitudes correlated with behavioral precision in this task. Finally, in Experiment 3 we manipulated the proportion of memory resources that should be allocated to lateral items by controlling the number of items presented laterally while manipulating overall load. Consistent with the first two experiments, we found that CDA amplitude tracked the proportion of memory resources allocated toward the lateral items in the memory array. When comparing across Experiments 2 and 3, we also found that CDA amplitudes were best predicted by a weighted sum of memory load and resources. This finding points to the CDA as a proxy of more than memory load alone, suggesting that this component may also reflect the total amount of memory resources allocated to lateral items.

One potential argument against the resource allocation interpretation is that instead of flexibly distributing resources across all items in the display, individuals alternated strategies *probabilistically* across trials: encoding higher probability items on most trials and lower probability items less often. If items were being encoded probabilistically, then one might expect that, on the majority of trials, participants would pay attention to and encode the higher probability items first. This would result in an earlier N2pc, consistent with previous findings in visual search tasks^32,33^. In contrast to this hypothesis, there was no difference in the timing of the N2pc across conditions in Experiment 2, suggesting that all items were being attended to at the same time, regardless of priority.

One interpretation of the probabilistic account is that participants alternate across trials between encoding high or low probability items; however, this interpretation can be ruled out based on the behavioral estimates of guess rates obtained from the two-component mixture model^34^ (see Supplementary Results). In Experiment 2, for example, if differences in CDA amplitudes were due to participants probabilistically encoding items, then the number of encoded items in the 75/25% condition should reflect this. That is to say, if participants encoded the two higher priority items on 75% of trials, and the two lower priority items on 25% of trials, this would lead to the same difference in CDA amplitude between conditions as was observed here, but with a total estimated number of encoded items of 2. However, the total number of encoded items in these conditions was 2.9, significantly greater than what would be predicted by the probabilistic encoding account.

Even if participants weren’t probabilistically encoding items, they may have been selectively encoding the higher priority items on the majority of trials, which could affect the CDA amplitude. However, this strategy cannot be ruled out based on estimates of the total number of items encoded, as increased guessing can also result from low resource allocation^18,35^. Indeed fluctuations in CDA amplitude across trials is likely a feature of all CDA measurements^36^, as changes in the number of items and amount of information encoded have been observed due to spontaneous fluctuations in attention^5,21,37,38^, and top-down strategies^39,40^. Thus, although the experiments presented here may include some measure of strategic differences in resource allocation across trials, resulting in some change in the number of stored items, *spontaneous* changes in resource allocation across trials may be an additional source of variance within past CDA studies that has been previously unidentified.

There are several implications that arise from these findings, such as the role of attentional enhancement in prioritization. In Experiment 2 we found that the N2pc, but not the P_D,_ tracked the priority of the lateralized items, while also predicting the precision of memory report. This suggests that when using feature-based cues, participants relied on up-weighting relevant information depending on their respective priorities, in comparison to down-weighting irrelevant information using active suppression. This is consistent with previous findings, which found that when given a pre-cue to up- or down-regulate memory encoding processes, participants could only up-regulate processing to benefit performance^3^. Additionally, it has been found that when using reward to prioritize items, only target selection is impacted and not distractor suppression^41^. This speaks to the importance of attentionally selecting and enhancing target information to VWM, instead of only inhibiting and filtering distractors. That is, although previous work has focused on the link between unnecessary memory storage of distractors and VWM capacity^42,43^, a full account of working memory performance should require a mechanism wherein resources are allocated amongst items when they are all relevant to the task^11^. The present results begin to point to potential mechanisms.

Finally, our results provide some information about the neuronal underpinnings of the CDA. The finding that the CDA follows a power-law when fit with a combination of resources and memory load is consistent with the saturation model of delay period activity proposed by Bays^17^. In this model, as input increases, neuronal activity also increases. However, as the input becomes increasingly large, it produces a consequent smaller increment in neuronal activity, saturating at some maximum level^17^. Although the power-law examined here tests a similar pattern, the present experiments did not test a large enough range of set sizes to delineate between capacity-limited models and limitless models. Moreover, it could be that prioritization is only possible within a limited range of stored items.

Our results do suggest that it is possible that different types of information can independently affect the CDA. Consistent with this interpretation, there are a few studies that have found memory precision can modulate the CDA independent from memory load^44,45^ (but see^46^). These previous findings support an account of VWM that encompasses both discrete item units, as well as continuous modulations in representational quality^47,48^. Consistent with this finding, a previous study observed independent effects of the number of items and complexity on the CDA at different channel pairs^49^. Thus, while the results of the experiments presented here cannot fully adjudicate between different models of VWM architecture (i.e., whether or not memory resources are tied to discrete units), they are consistent with the idea that how resources are allocated should be considered in addition to overall load in neural and behavioral models of VWM.

## Methods

### Participants

Informed, written consent was obtained from all participants. Procedures were approved by and conducted in accordance with the Brock University Bioscience ethics review board. We aimed for a sample size of 20 participants^15,16,50^ (right handed, normal-color vision, no history of mental illness). To reach these targets, a total of 30 participants were run in Experiment 1, 33 in Experiment 2, and 28 in Experiment 3. Distinct samples (N = 20) were used for each experiment (Exp 1: *M*_age_ = 22.0, *SD*_age_ = 3.0, 10 male; Exp 2: *M*_age_ = 22.6, *SD*_age_ = 4.2, 9 male; Exp 3: *M*_age_ = 21.6, *SD*_age_ = 3.9, 3 male).

### Stimuli and Procedures

All tasks were completed on a Windows PC with a 41-cm NEC MultiSync LCD 2090UXi computer monitor (1600 × 1200 pixels, 60 Hz refresh rate). Stimuli were rendered using Psychopy v1.90.3 (Peirce, 2007) and presented on a grey background (RGB = 128 128 128) with a central fixation dot (radius of 0.3°). Viewing distance was approximately 57 cm. In all experiments, participants first completed a standard change detection task^51^. These data are not included.

The colors for the squares in the continuous report VWM tasks were chosen pseudo-randomly from a 360-degree isoluminant color wheel (CIE L*a*b* color space, [L = 70, a = −6, b = 14, radius = 49]), which was calibrated to the testing monitor. Memory stimuli colors were separated by at least 30 degrees on the color wheel.

#### Experiment 1

At the beginning of each block, participants were instructed on cue-validity, both with on-screen and verbal instructions: In the 1-Cue/100%-Valid condition, the one cued item was always probed; In the 4-Cues/100%-Valid condition, the probed item was randomly selected from all four items; In the 1-Cue/50%-Valid condition, the cued item was probed on 50% of trials, un-cued items were probed on the remaining trials. Each trial began with an arrow indicating which side of the screen was task relevant (3° tall, 200 ms), followed by a jittered fixation interval (200 – 500 ms). The memory array (150 ms) consisted of four squares on both sides of the screen (1° x 1°, 4° from fixation) along with horizontal spatial line cues (2° long x 3 pixels wide, 2° from fixation). After a delay (900 ms), participants reported the color of the bolded square (line width of 3 pixels vs. 1 pixel) from the color wheel (diameter of 7°) with the mouse. There were 240 trials in both the 1-Cue/100%-Valid and 4-Cues/100%-Valid Conditions, and 480 in the 1-Cue/50%-Valid condition, split equally between both sides of the screen (total 960). One participant’s data consisted of only 840 trials due to a recording error.

#### Experiment 2

In both Experiments 2 and 3 participants first completed a subjective luminance-matching task, which was used to create the placeholder colors (see Supplementary Methods).

Each trial began with a feature-based cue (i.e. *100% Circle*; 1,000 ms, 1.5° tall) followed by a jittered interval (500 – 1,000 ms). Next, eight shapes were presented in clusters of two at each cardinal position 3° from fixation (to center of shape cluster) and 1.2° apart (vertical center to center; 200 ms). There were always two colored squares (1° × 1°) and two colored circles (diameter of 1°). The remaining four items were filled with the subjectively luminance-matched grey and were always the un-cued shape. Shapes were presented in all possible position configurations equally (16 unique positions). After a delay (900 ms), participants made a response to the probed shape on the color wheel with the mouse (pseudo-randomly chosen from top or bottom shape in cluster). There were a total of four conditions defined by the probability that the lateral shapes would be probed: 100%, 75%, 25%, and 0%. Participants completed a total of 816 trials (100% lateral: 200, 0% lateral: 200, 75% lateral: 208, 25% lateral: 208). One participant completed 806 trials due to a programming error, and another completed 807 trials due to an interruption to the recording session.

#### Experiment 3

Participants were instructed that all items were equally likely to be probed. Each trial began with a jittered ITI (500 – 1,500 ms) followed by the memory array (200 ms). Then, after a delay (900 ms), participants made a response to the probed shape on the color wheel with the mouse. There were three conditions: 1) Load 4 with three colored squares presented in a vertical cluster to the left or right of fixation (1° × 1°, 1.2° apart center-to-center, 3° from fixation) and one colored square presented vertically. 2) Load 4 with one square presented laterally and three squares on the vertical. 3) Load 6 with three squares presented laterally and 3 vertically. Participants were given feedback after their response (800 ms), where ‘Correct’ was considered within 40° of the target color. Participants completed a total of 900 trials, 300 of each condition.

### EEG Recording and Pre-Processing

All EEG pre-processing was done in MATLAB with the EEGLAB^52^ (Version 14.0.0b), and ERPLAB^53^ (Version 6.1.2) toolboxes. EEG was DC recorded at a 512 Hz sampling rate from a 64 Ag/AgCl electrode cap placed at the standard 10-20 sites^54^. The signal was online referenced to the common mode sense (CMS) and the driven right leg (DRL) electrodes. Data were re-referenced off-line to the average of the mastoids, baseline corrected to −200 ms before memory array onset, and filtered with a 40-Hz low-pass and 0.1-Hz high-pass Butterworth filter (slope: 12dB/octave). Data were epoched between −200 and 1,050 ms (Experiment 1) or −200 and 1,100 ms (Experiment 2), time-locked to the memory array.

### Artifact rejection

Horizontal electro-oculogram (HEOG) was recorded from bipolar external electrodes placed laterally beside the eyes. Vertical electro-oculogram (VEOG) was recorded as the difference between external electrodes placed below the eyes and FP1 or FP2. Trials with VEOG activity > ±80 μV or HEOG activity > ±32 μV between stimuli onset and the end of the trial were removed, as were trials in which the voltage over posterior channels (P1/2, P3/4, P5/6, P7/8, P9/10, PO3/O4, PO8/O7, and O1/2) was > ± 100 μV. Participants with more than 35% of trials rejected were replaced. An average of 21.4% of trials were rejected in Experiment 1, (*SD* = 10.4%), 11.6% in Experiment 2 (*SD* = 7.8%), and 13.3% in Experiment 3 (*SD* = 10.1%). Across studies, each participant had more than 100 trials in each condition bin.

In Experiments 2 and 3 we also replaced participants whose average residual HEOG activity was greater than ± 4 μV between memory array onset and the end of the epoch. Resulting in absolute residual HEOG values of 1.61 μV (*SD* = 1.02 μV) in Experiment 2 and 1.61 μV (*SD* = 1.00 μV) in Experiment 3. The total number of replaced participants was 10 in Experiment 1, 12 in Experiment 2, and 8 in Experiment 3.

### Data Analysis

#### Behavioral data

Performance was assessed using the trial-by-trial raw response error (i.e., the difference in degrees between the color of the probed item and the participant’s response). Lower values reflect more precise responding. Data from all experiments were fit to a power-law function:

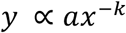

Bayesian information criterion (BIC) values were computed to compare model fits. Raw error values were calculated using custom scripts in MATLAB. Goodness of fit was computed using nonlinear least squares regression in MATLAB’s Curve-Fitting Toolbox using a bisquare robust fitting procedure with the group data averaged across conditions.

#### ERP data

Difference waves were calculated at electrode pairs: P3/4, P7/8, PO7/O8, PO3/O4, and O1/2^55^. Across all experiments, there was no significant Condition x Channel interaction for any of the ERP components (*F*s < 2.24, *p*s > .055, η^2^_p_s < .11). Therefore, we averaged activity across these electrode sites for all ERP measurements.

Repeated-measures ANOVAs were used along with follow-up linear contrasts and fits where reported. Where violations of sphericity exist, Greenhouse-Geisser corrected degrees of freedom and *p* values are reported. Two-tailed t-tests were Bonferroni-corrected only where a priori hypotheses were not present. Bayesian repeated-measures ANOVAs and post-hoc tests are reported where applicable (r scale prior width of 0.5, default Cauchy prior centered on 0, 10,000 Monte Carlo samples). Statistical analyses were completed using JASP Version 0.8.4^56^ and MATLAB R2017a.

In all experiments, the CDA was measured as the mean amplitude during the delay period from 400 ms post-stimuli offset to the end of the trial^36,57^. An N2pc was only observed in Experiment 2 and was measured as the mean amplitude from 200 – 300 ms post-stimuli onset^32,58–60^. We also measured the negative 50% fractional peak latency of the N2pc between 200 – 300 ms^61^. In Experiment 2, we measured the P_D_ as the positively signed area from 250 – 400 ms^62,63^. We used a nonparametric permutation approach to determine the presence of the P_D_.

The *p* values for the permutation tests were estimated using the following formula with 1,000 permutations^64^:

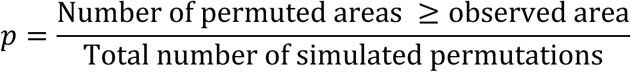

#### Modelling CDA data

In Experiments 2 and 3, we fitted the CDA amplitudes to a power-law model. The weighted-product values were calculated by the following formula:

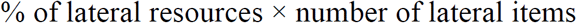

Model fits were completed using the Curve Fitting Toolbox in MATLAB and custom MATLAB scripts to calculate BIC values to compare model fits.

## Data Availability

The datasets generated or analysed during the present study are unavailable due to the absence of consent.

## Code Availability

Psychopy task scripts, R code, and MATLAB code used to pre-process EEG data as well as to fit models are available online at: https://osf.io/r73c5/

## Acknowledgements

This work was supported by NSERC grants (#435945 and #458707) awarded to S.M.E., and NSERC PGS-D scholarships awarded to C.S. and H.A.L. We thank Joey Capozza (experiment programming/data collection), Thomas Nelson and Kevin MacDonald (experiment programming), and Brenda de Wit (data collection).

## Author Contributions

C.S., H.A.L., N.A., and S.M.E conceived of the experimental designs. C.S., H.A.L., and S.M.E. analyzed the data; C.S., H.A.L., and B.D. collected the data. C.S. and S.M.E. took the lead in preparing the manuscript. All authors discussed the interpretation of results and contributed to the final manuscript.

