## Supplemental Methods for "Electrophysiological correlates of the flexible allocation of visual working memory resources"

**memory resources**

Christine Salahub^1*^, Holly A. Lockhart^1^, Blaire Dube^2^, Naseem Al-Aidroos^3^, and Stephen M. Emrich^1^

^1^Department of Psychology, Brock University

St. Catharines, Ontario, Canada

^2^Department of Psychology, The Ohio State University

Columbus, Ohio, USA

^3^ Department of Psychology, University of Guelph

Guelph, Ontario, Canada

**Supplementary Methods.**

**Luminance-matching task**

This task was used in Experiments 2 and 3 to obtain an individual grey color that was subjectively matched in brightness to the color wheel colors. This grey was used in the main experiment as the fill color for the placeholder shapes to visually balance the display. In this task, participants were presented with a screen that was vertically divided in half with a fixation point (0.3° x 0.3°) in the middle. The left half of the screen was filled with a color from the color wheel and the right half was filled with grey. Twelve colors were selected from the color wheel (in 30° steps). Each color was presented four times: twice each with a grey that was ~5 cd/m^2^ greater or less than the average brightness of the 12 colors (~25.1 cd/m^2^). Participants were instructed to match the brightness of the grey half of the screen to that of the color using the left and right arrow keys. Each press of the right arrow key increased the lightness of the grey by 0.01 units closer to white, and the left arrow key decreased the lightness by 0.01 units closer to black (-1 to 1 RGB color space where [0, 0, 0] is grey). To make their final response, participants pressed the enter key. For each individual participant we used their averaged grey luminance across all 48 ratings as the fill color for the placeholders in Experiment 2 (*M* = [-.02 -.02 -.02] or ~35.6 cd/m^2^, *SD* = .087) and Experiment 3 (*M* = [-.047 -.047 -.047] or ~29.3 cd/m^2^, *SD* = .041).

**Supplementary Results.**

**Standard mixture model estimates of visual working memory capacity**

Behavioral data from all three experiments were analyzed using the two-component standard mixture model (Zhang & Luck, 2008). This model provides two parameter estimates per experimental condition: P_m_, the probability that the probed item was encoded and stored in memory; and standard deviation (SD), which reflects the precision of the memory representation. Here, we focus on the effect of experimental condition on P_m_ as a way to examine whether participants were probabilistically encoding items into memory in each experiment. To calculate the number of items held in memory, we used the following calculation:

$$P_{m}\times Load$$

Where Load refers to the total number of items that should be held in memory if performing the task correctly. P_m_ can also be calculated as (1 – Guess Rate).

**Experiment 1.**

In this experiment, we defined Load as 1 when one item was cued with 100% validity and 4 when 1 item was cued at 50% validity or when all 4 items were cued. A repeated measured ANOVA was conducted on the estimates of memory load for each condition. It was found that there was a main effect of overall Condition, *F*(1.39, 26.40) = 87.51, *p* < .001, η^2^_p_ = 0.82, BF_10_ = 1.42 x 10^14^. More items were held in memory when all 4 items were cued with 100% validity (*M* = 1.98, *SD* = 0.58), than when only 1 item was cued with 100% validity (*M* = 0.97, *SD* = 0.02), *t*(19) = -7.78, *p_bonf_* < .001, d = 1.74, BF_10_ = 6.29 x 10^4^. However, participants held more items in memory when 1 item was cued with 50% validity (*M* = 2.23, *SD* = 0.35) than when all 4 items were cued, *t*(19) = 2.86, *p_bonf_* = .03, d = 0.64, BF_10_ = 5.14, and more than when 1 item was cued with 100% validity, *t*(19) = 16.32, *p_bonf_* < .001, d = 3.65, BF_10_ = 5.65 x 10^9^ (see Table S1 for a summary of the guess rates and memory load estimates). Contradictory to the pattern of CDA amplitudes, these findings suggest that more items were being held in memory when 1 item was cued at 50% validity than when all 4 items were cued with 100% validity. This suggests that in this condition, participants were not strategically holding only the highest probability item in memory on the majority of trials.

**Experiment 2.**

Here, we focused on P_m_ in the 75/25% condition to determine whether participants were probabilistically encoding the memory items. If participants were encoding the two higher priority items on 75% of trials, and the two lower priority items on the remaining 25% of trials, this would result in a pattern of CDA amplitudes similar to what was observed in the current study. However, this probabilistic pattern of responding should be reflected in behavioral estimates of memory load (i.e. P_m_ * Set Size). That is to say, if participants were encoding the two 75% likely items on 75% of trials, an average of 75 * 2 = 1.5 items should be encoded into memory. Adding this to the number of items encoded on the remaining 25% of trials results in a total average of 1.5 + 0.5 = 2 items. Therefore, if participants were probabilistically encoding items into memory, the number of items in memory in the 75/25% condition should not be significantly different from 2. Similarly, if participants were only holding the higher priority items in memory, we would also expect 2 items to be held in memory on average. However, if participants were instead using flexible resource allocation amongst all items in the memory display, then the number of items in memory should be greater than 2.

Although we focus our analysis on 75/25% condition, all estimates per Condition are presented in Table S2 (excluding the 0% priority condition). Collapsed across probed item priority, an average number of 2.92 items were held in memory in the 75/25% condition (*SD* = 0.59). This is significantly different from 2, *t*(19) = 7.05, *p* < .001, d = 1.58, BF_10_ = 1.74 x 10^4^. When the same analysis was applied to each probed condition separately, it resulted in a number of poor model fits. Nevertheless, the total number of items encoded in memory (*M* = 2.33, *SD* = 0.64) was significantly greater than the 2 predicted by the probabilistic account, *t*(19) = 2.33, *p* = .031, BF_10_ = 2.04. These findings suggest that participants were not probabilistically encoding the items in the 75/25% condition, and were instead using flexible resource allocation to encode all items in the display.

**Experiment 3.**

Guess rates and estimates of the number of items held in memory for Experiment 3 are presented in Table S3. A paired samples t-test comparing the number of items held in memory in the Load 4 condition (collapsed across which item was probed) and the Load 6 condition, showed that more items were stored in memory at Load 4 (*M* = 1.90, *SD* = 0.47) than in the Load 6 condition (*M* = 1.63, *SD* = 0.71), *t*(19) = 2.99, *p* = .007, d = .67, BF_10_ = 6.58. Using the same logic as presented in Experiment 2, in the Load 4 condition if participants were holding 3 items in memory on 75% of the trials, then they would on average store 2.25 items in memory. This is then added to the 25% of trials where 1 item was remembered, resulting in an average of 2.25 + 0.25 = 2.5 items for the Load 4 condition. Similarly, for the Load 6 condition, the probabilistic account would predict that 50% of the time only three items are being remembered, resulting in an average of 3 items. Thus, although the total number of encoded items was less than what would be predicted, there were a greater number of items encoded in the Load 4 condition, contrary to the probabilistic account.

Table S1

*Guess Rates and Memory Load Estimates by Condition in Experiment 1*

Condition	*M(SD)_Guess_	M(SD)_Load_*

1 Cue, 100% Valid	0.03 (0.02)	0.97 (0.02)

1 Cue, 50% Valid	0.44 (0.09)	2.22 (0.35)

Cued Item Probed	0.09 (0.08)	0.91 (0.08)

Non-Cued Item Probed	0.68 (0.27)	0.96 (0.81)

4 Cues, 100% Valid	0.51 (0.15)	1.98 (0.58)

*Note.* N = 20.

Table S2

*Guess Rates and Memory Load Estimates by Condition in Experiment 2*

Condition	*M(SD)_Guess_	M(SD)_Load_*

100/0%	0.18 (0.10)	1.64 (0.20)

75/25%	0.27 (0.15)	2.92 (0.59)

75% Probed	0.24 (0.11)	1.52 (0.23)

25% Probed	0.59 (0.34)	0.81 (0.69)

*Note.* N = 20.

Table S3

*Guess Rates and Memory Load Estimates by Condition in Experiment 3*

Condition	*M(SD)_Guess_	M(SD)_Load_*

Load 4	0.52 (0.12)	1.90 (0.47)

25% Probed	0.24 (0.14)	0.76 (0.14)

75% Probed	0.61 (0.15)	1.16 (0.44)

Load 6	0.73 (0.12)	1.63 (0.71)

*Note.* N = 20.
